# Orientation of Directional Deep Brain Stimulation Leads on CT: Resolving the Ambiguity

**DOI:** 10.1101/2020.09.16.298653

**Authors:** Rebecca Kurtev-Rittstieg, Stefan Achatz, Amir Nourinia, Stephan Mittermeyer

**Author notes:** Corresponding Author: Rebecca Kurtev-Rittstieg, R&D Functional and Stereotactic Neurosurgery, Brainlab AG, Olof-Palme-Str. 9, 81829 Munich, Germany.

## Abstract

While directional deep brain stimulation (DBS) shows promising clinical effects by providing a new degree of freedom in programming, precise knowledge of the lead position and orientation is necessary to mitigate the resulting increased complexity. Two methods for orientation assessment based on postoperative CT imaging have become available, but neither of them is currently able to resolve the respective 180° artifact symmetry. Both rely on information about the intended orientation and assume that a deviation of more than ± 90° is very unlikely. Our aim was to develop an enhanced algorithm capable of detecting asymmetries in the CT data and to thus eliminate the need for user interaction. Two different approaches are presented: one based on the lead marker’s center of mass (COM) and one based on asymmetric sampling of the marker’s intensity profile (ASM). Both were tested on a total of 98 scans of 2 lead phantoms, resulting in 165 measurements with a large variety of lead implantation and orientation angles. The 180° ambiguity was correctly resolved in 99.4% of cases by COM and in 96.4% of cases by ASM. These results demonstrate the substantial and currently unused asymmetry in CT and the potential for a truly automated workflow.

## Introduction

Directional deep brain stimulation (DBS) capable of asymmetrically shaping the electrical field has shown to increase side-effect thresholds and improve the therapeutic window when compared to conventional DBS [1, 2]. These promising clinical effects, however, come at the cost of increased complexity. In order to facilitate programming, stimulation parameters need to be related to patient-specific neuroanatomy, which requires not only knowing the lead’s position but also its orientation [3]. Surgical control of the latter has proven challenging, and significant deviations from the intended orientation have been demonstrated [4]. Therefore, various methods using postoperative imaging have been proposed, such as stereotactic X-ray, flat-panel computed tomography (fpCT), 3D rotational fluoroscopy (RF) and standard CT [3, 5–7]. The CT-based approach has become the most widely adopted, as CT is already part of the standard postoperative workflow in most centers. Two options for such automatic orientation assessment are currently available, based on the same characteristic artifacts: the DiODe algorithm as integrated into the open-source Lead-DBS toolbox [3, 4] and the commercial software package Elements® Lead Localization (Brainlab AG, Munich, Germany). However, the underlying artifacts exhibit a 180° symmetry (Fig. 1), and neither method is currently able to resolve this ambiguity. Both rely on the intended orientation and assume that a deviation of more than ± 90° is very unlikely. If such a deviation occurs in clinical practice, it can lead to a wrong hypothesis for image-guided programming or errors in the analysis of stimulation effects.

**Fig. 1.**
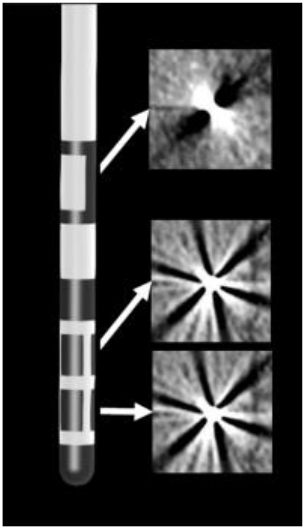
Illustration of the Boston Scientific Vercise Cartesia™ directional lead (left) and its characteristic artifact patterns (right), which exhibit a strong 180° symmetry at standard clinical CT resolution. The lead features a 1-3-3-1 design with two levels of segmented electrodes in the center, enclosed by one cylindrical electrode at each end, and a radiopaque orientation marker located dorsally to the electrodes with a 2 mm marker band spanning 140°.

In this study, we aimed to develop an enhanced algorithm capable of detecting slight asymmetries in the respective CT artifacts, thus providing a fully automated workflow. Two approaches are introduced and tested on phantoms: one based on the lead marker’s center of mass (COM) and one based on asymmetric sampling of its intensity profile (ASM).

## Methods/Design

### Phantom Design

Two Vercise Cartesia™ DBS leads (Boston Scientific, Marlborough, MA, USA) were each embedded into an acrylic rod by a precision mechanic and three metal stylets were added at the ends parallel to the lead axis. MicroCTs with 70 µm resolution (MITOS GmbH, Garching bei München, Germany) were acquired to determine each lead’s ground truth orientation with respect to the reference stylets (Fig. 2a). Two setups were created; a water-filled box with 3D printed custom turntables (Fig. 2b) and a water-filled anthropomorphic skull (TruePhantoms Solutions Inc, Windsor, ON, Canada, Fig. 2c). The box enabled imaging at precise polar and orientation angles, where polar angle refers to the angle between lead axis and CT slice normal vector. The skull featured realistic adult bone thickness and Hounsfield unit (HU) values as well as custom inserts based on standard coordinates for subthalamic nucleus DBS.

**Fig. 2.**
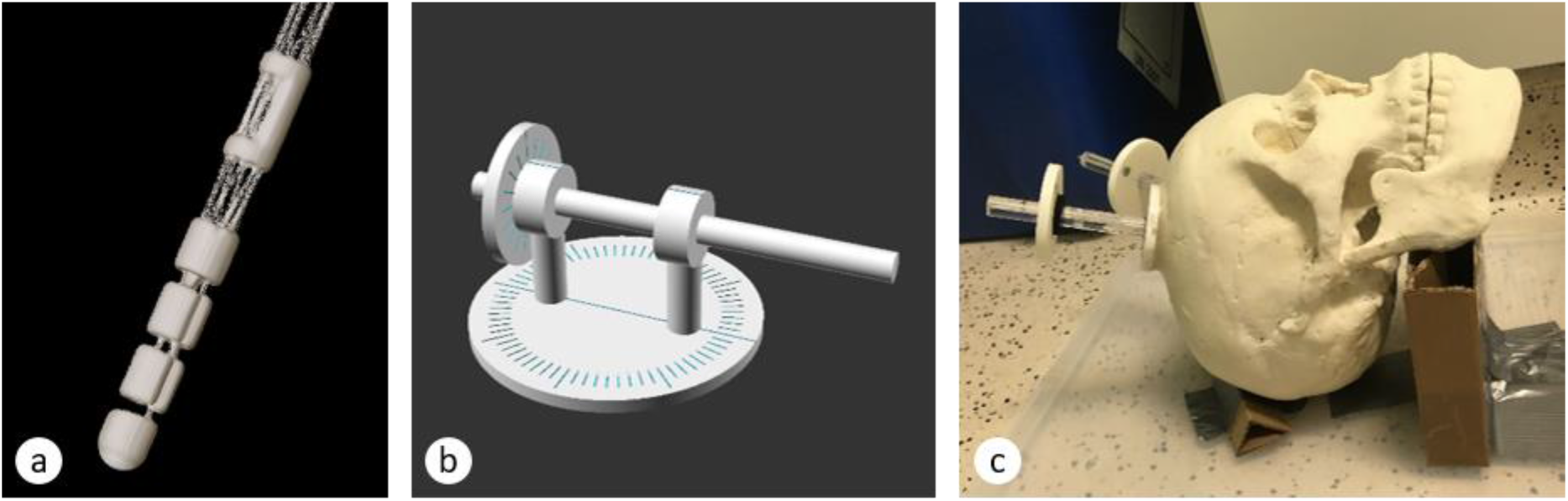
Illustration of the phantom setups. **(a)** 3D reconstruction of a microCT of a Boston Scientific Vercise Cartesia™ directional lead embedded in an acrylic rod at 70 µm resolution. Reference stylets cut off for visualization purposes. **(b)** Turntable phantom design with an acrylic rod allowing for water box scans at precise polar and orientation angles. **(c)** Anthropomorphic skull with both acrylic rod phantoms reflecting standard DBS implantation positions.

### Data acquisition

A total of 98 CT scans were acquired using various scanners and standard DBS protocols, featuring both axial and helical acquisitions and a slice thickness ≤ 1 mm. All scans were performed with 0° tilt and manufacturer-specific metal artifact reduction turned off. Details are provided in Table 1.

**Table 1.** Overview of the used CT scanners and their respective imaging parameters.

| Manufacturer | Model name | Slice thickness (mm) | Axial / helical mode | Pitch factor for helical mode | Voltage (kV) | Exposure (mA) |
| --- | --- | --- | --- | --- | --- | --- |
| Mobius Imaging | AIRO | 1.0 | axial | NA | 120 | 169 |
| Philips | Brilliance 64 | 0.625 / 0.8 | axial / helical | 0.31 | 120 | 300 / 103 |
| Philips | Ingenuity CT | 0.8 | axial / helical | 0.3 | 120 | 350 / 99 |
| Philips | Ingenuity Flex | 0.75 / 0.8 | axial / helical | 0.44 | 120 | 400 / 105 |
| Siemens | Emotion 16 | 0.6 / 0.75 / 1.0 | axial / helical | 1.05 | 130 | 120 / 180 / 300 / 154 |
| Siemens | SOMATOM Definition AS | 1.0 | axial / helical | 0.55 | 120 | 209 |

The lead polar angles in the water box setup (n = 91) ranged from 0.1° to 49.6° (median 27.2° ± 14.7°) and those in the skull setup (n = 76) from 23.4° to 44.6° (median 35.6° ± 6.3°). The orientation angles as transferred from microCT via rigid registration ranged from −180° to 179.9° to DICOM anterior in the water box setup (median −27.5° ± 95.2°) and from −174.5° to 175.7° in the skull setup (median 11.2° ± 96.9°). The respective distributions are illustrated in Figure 3.

**Fig. 3.**
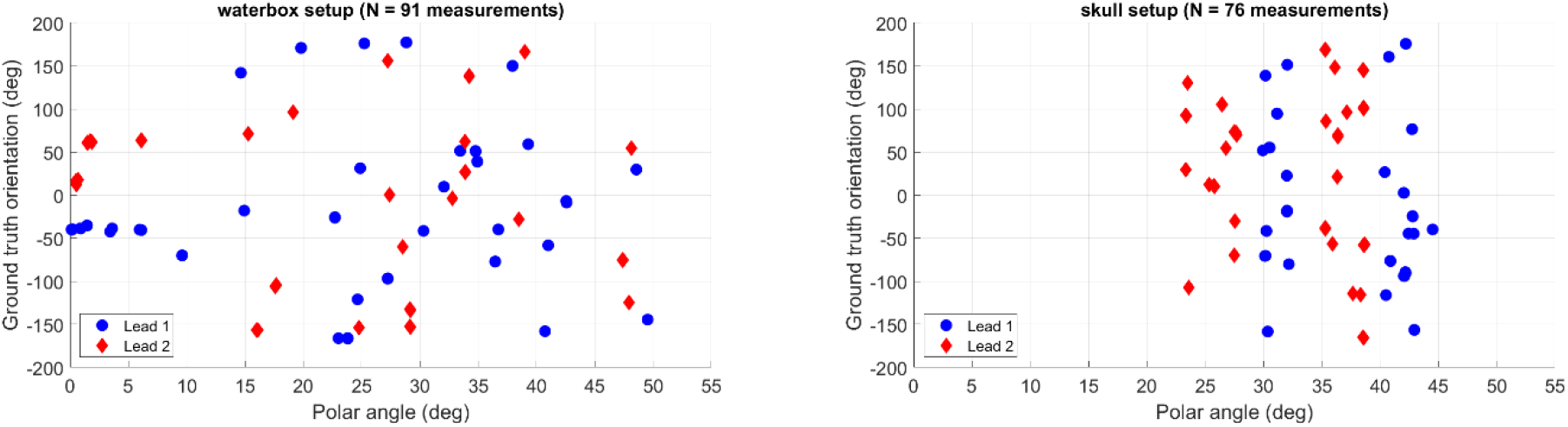
Scatter plot of the analyzed lead polar angles and orientation angles.

### Algorithms

Both approaches introduced here build upon the algorithm available in Elements® Lead Localization (Brainlab AG, Munich, Germany), which is referred to as L-Orient in the following. In a first step, L-Orient models the expected two-streak artifact pattern for all CT slices intersecting the marker band. Intensity values are extracted at varying distances from the lead-slice intersection point and averaged (Fig. 4a). This is repeated in 360 increments of 1° to reflect a full lead rotation and aggregated into an intensity profile (Fig. 4d). Within the optimal slice as defined by the global minimum, two local minima from the marker are expected. If this is fulfilled, an analogous analysis is conducted for the segmented electrodes by modeling the six-streak artifact pattern as in the DiODe approach (Fig. 4c). L-Orient then searches for a local minimum within ± 20° of each of the two possible marker orientation angles and within ± 60° of the intended orientation.

**Fig. 4.**
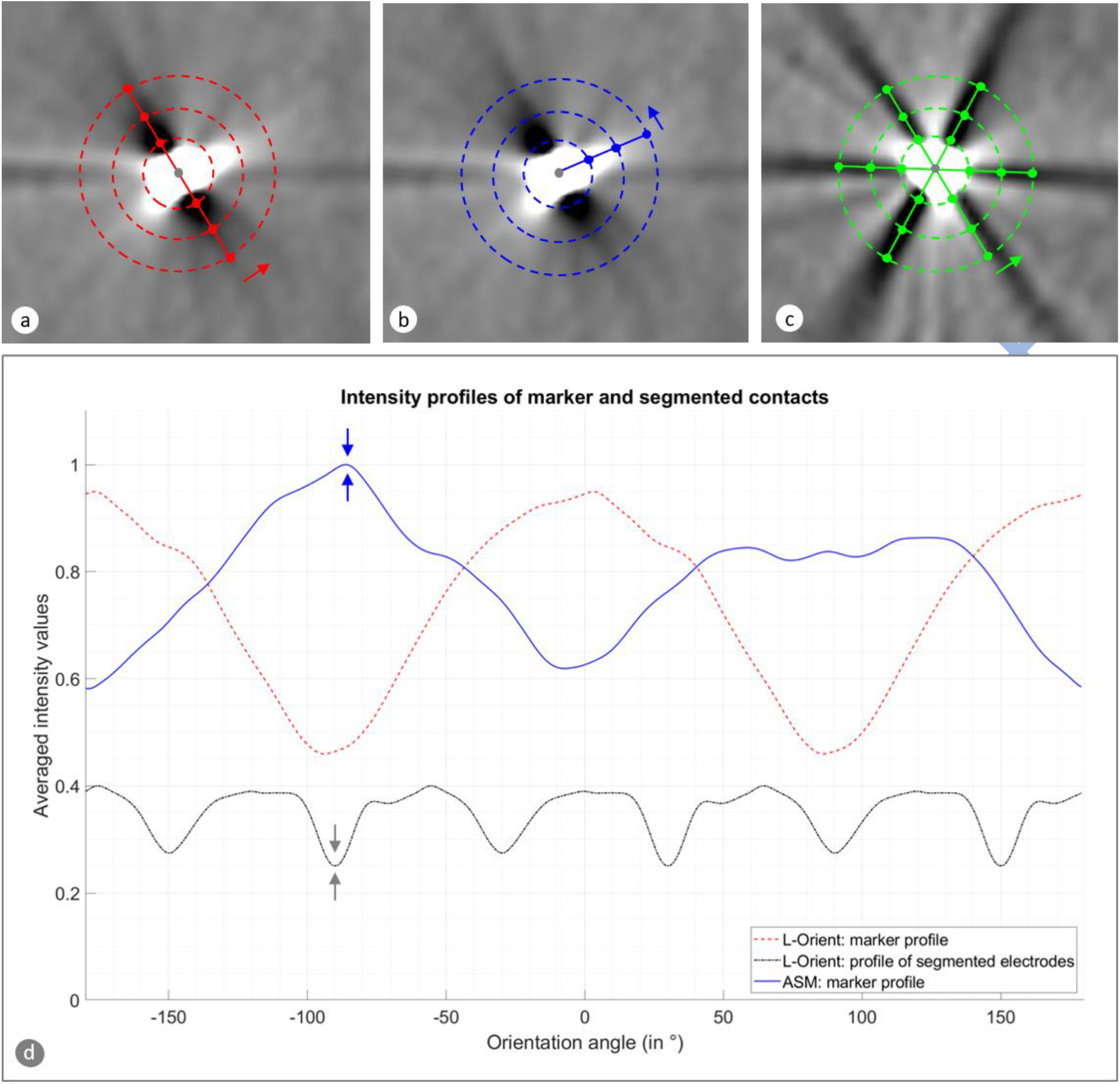
Representation of the artifact analysis on selected CT slices and the resulting intensity profiles for a directional lead with polar angle 39.3°. **(a)** CT slice at marker level with model of the expected two-streak artifact pattern as implemented in L-Orient. Sampling points visualized at 2 mm, 4 mm, and 6 mm distance to the lead-slice intersection point. **(b)** CT slice at marker level with asymmetric sampling as in the ASM approach. **(c)** CT slice at the level of segmented electrodes with a model of the expected six-streak artifact pattern, as used in all approaches. **(d)** Intensity profiles resulting from the sampling illustrated in (a)-(c), normalized for visualization purposes. The L-Orient marker intensity profile (red) exhibits two minima coinciding closely with local minima of the intensity profile of the segmented electrodes (black), but these two possible solutions of −88° and 92° to anterior for the lead orientation cannot be distinguished well. However, a global maximum can be found for the ASM marker intensity profile (blue), which leads to the correct solution of in this case −88°.

### Marker center of mass (COM)

This approach hypothesizes that the 140° marker band causes hyperintense CT artifacts which are slightly asymmetric with respect to the lead axis. The region around the marker is therefore assessed by re-sampling slices perpendicular to its axis with an in-plane resolution of 0.1 mm and a slice thickness of 0.2 mm. A threshold of 2000 HU is applied to segment the marker volume and calculate its center of mass. The 3D distance vector from the lead axis to this center of mass is then used to determine the most likely lead orientation from the two potential L-Orient solutions.

### Asymmetric sampling of the marker intensity profile (ASM)

This approach is based on the hypothesis that the bright ellipsoidal region around the intersection of the marker with the CT slice should be asymmetric in the direction of the marker band. Analogous to L-Orient, the expected artifact pattern is modeled by a line through the lead-slice intersection point, but here the sampling points are placed along half of it instead of symmetrically into both directions. Intensity values are extracted and averaged (Fig. 4b). This is repeated in 1° increments and aggregated into a curve. Out of the two resulting local maxima, one is expected to be more pronounced due to the asymmetric nature of the marker (Fig. 4d).

### Statistical analysis

Both COM and ASM were assessed by how many cases they classified correctly, defined as a determined marker angle within ± 90° of the ground truth orientation. In terms of accuracy, we focused on the corresponding L-Orient results, which are based on the more precise segmented electrode streak artifacts. The Mann-Whitney U test was used to assess differences in the determined orientation angles grouped for the two lead phantoms and p ≤ 0.05 was used as significance level. Statistical analysis was performed using MATLAB version R2020a (MathWorks, Natick, Massachusetts, USA).

## Results

In our analysis of L-Orient results, box plots showed a characteristic drift over time when grouping by phantom and scan date. Therefore, additional microCTs were acquired. These revealed that lead 1 had rotated inside the acrylic rod by 10° over 23 weeks and lead 2 by 23° over 11 weeks. While the exact cause remains unclear, multiple factors may have contributed, such as suboptimal fixation within the rods, tension introduced during insertion, or impact during transport. As it was not possible to determine when exactly each lead rotation took place, we utilized the respective mean orientations as ground truth.

Significant (P = 1.4·10^−14^) differences between the two phantoms were found for the orientations detected by L-Orient, with a mean deviation of −0.4° ± 4.6° (range: −9.8° to 7.2°) to ground truth for lead 1 and 6.5° ± 5.2° (range: −8.0° to 26.1°) for lead 2. To correct for these imperfections, we subtracted the respective group mean from the individual measurements for each phantom analogous to Sitz et al. [5]. This resulted in a mean deviation of 0.0° ± 4.6° (range: −9.4° to 7.6°) for lead 1 and 0.0° ± 5.2° (range: −14.5° to 19.6°) for lead 2 (Fig. 5). In 2 out of the 167 analyzed cases, L-Orient could not determine the orientation, as it detected four minima in the marker intensity profile instead of two. For the remaining 165 lead measurements, the COM method succeeded in correctly resolving the 180° ambiguity in 164 cases, corresponding to a rate of 99.4%. The ASM algorithm correctly classified 159 out of 165 and hence 96.4% of cases. The distributions of correctly and falsely classified lead orientations are illustrated in Fig. 6.

**Fig. 5.**
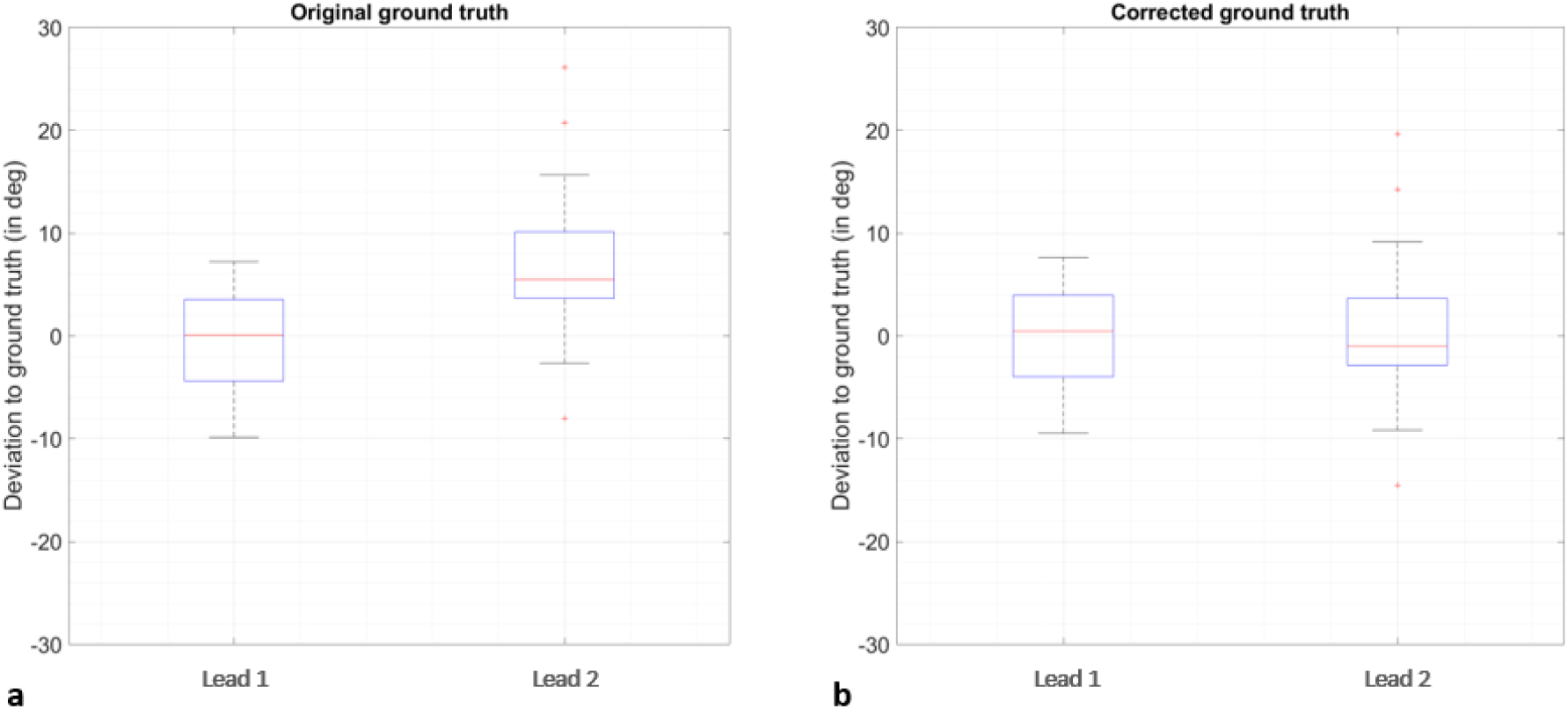
Lead orientation angles determined by L-Orient with respect to the ground truth defined by microCT shown as box plots for the two lead phantoms **(a)** before and **(b)** after correction for imperfections. Measurements for water box and skull setup are combined.

**Fig. 6.**
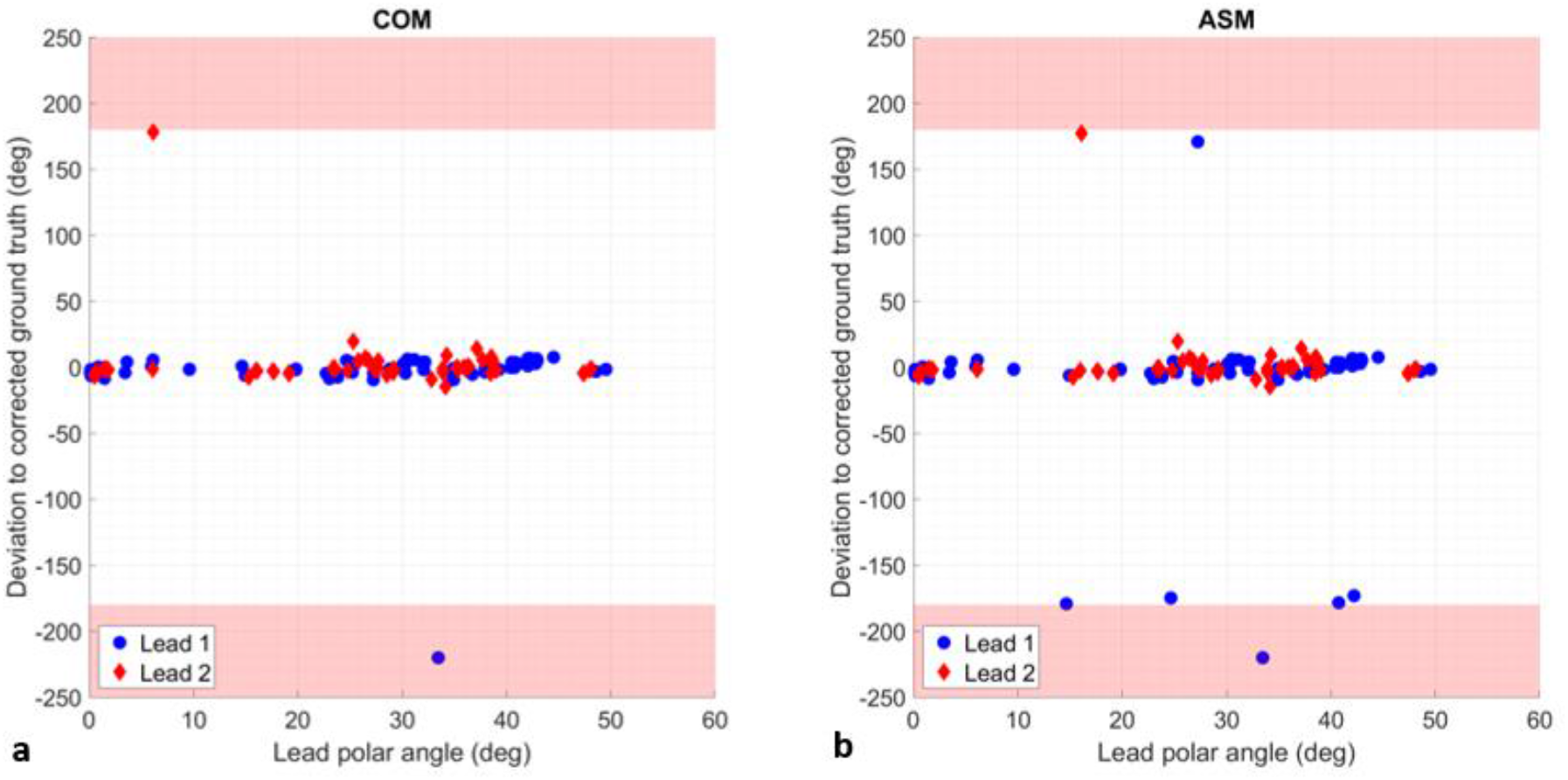
Distribution of determined lead orientation angles with respect to ground truth and polar angle. Cases where L-Orient could not determine a lead orientation were set to −220° for illustration purposes, and the angle range outside of [-180°, 180°] is highlighted in red. **(a)** COM method. **(b)** ASM method.

## Discussion/Conclusion

Both currently available methods for CT-based assessment of directional DBS lead orientation rely on the intended orientation and assume that a deviation of more than ± 90° is very unlikely. However, if such a deviation occurs, it can lead to a wrong hypothesis for image-guided programming. While this would not pose an immediate risk to the patient but result in the currently established trial-and-error-programming, it would create an unnecessary burden for the overall workflow.

We therefore aimed to develop an enhanced algorithm capable of detecting asymmetries in the respective lead artifacts. The COM and ASM approaches presented here were able to resolve the supposed 180° ambiguity in almost all cases, for a variety of scanners, lead implantation angles and orientations. No dependence on the lead polar angle was observed. It should be noted though that L-Orient is restricted to polar angles ≤ 50° in order to robustly evaluate the 2 mm marker band on standard postoperative DBS protocols with CT slice thickness ≤ 1 mm. DiODe was tested for a substantially smaller slice thickness of 0.8 mm and could thus cope with polar angles up to 60° [3].

Future investigations will focus on improvements in the manufacturing of lead phantoms, a systematical analysis of the dependence of the COM and ASM method on implementation parameters, and a possible combination of both.

Even though this study represents a proof of concept and further validation on patient data would be necessary to ensure the clinical reliability required for implementation in a medical device, our results demonstrate the potential for a truly automated and robust workflow. The latter is crucial in mitigating the increased complexity of directional DBS through patient-specific visualization, particularly when integrating volume of tissue activated (VTA) models for guiding programming and for better understanding the clinical effects of stimulation.

## Statements

### Author Contributions

S.A. and S.M. designed the research project. S.A. and A.N. executed the research project. R.K.-R. and S.A. analyzed the data and wrote the article. All authors were involved in the discussion of the results and commented on the article.

